# Activity Map and Transition Pathways of G Protein Coupled Receptor Revealed by Machine Learning

**DOI:** 10.1101/2022.12.20.521237

**Authors:** Parisa Mollaei, Amir Barati Farimani

## Abstract

Approximately, one-third of all FDA-approved drugs target G protein-coupled receptors (GPCRs). However, more knowledge of protein structure-activity correlation is required to improve the efficacy of the drugs targeting GPCRs. In this study, we developed a machine learning (ML) model to predict activation state and activity level of the receptors with high prediction accuracy. Furthermore, we applied this model to thousands of molecular dynamics trajectories to correlate residue-level conformational changes of a GPCR to its activity level. Finally, the most probable transition pathway between activation states of a receptor can be identified by using the state-activity information. In addition, with this model, we can associate the contribution of each amino acid to the activation process. Using this method we will be able to design drugs that mainly target principal amino acids driving the transition between activation states of GPCRs. Our advanced method is generalizable to all GPCR classes and provides mechanistic insight into the activation mechanism in the receptors.

## Introduction

The G protein-coupled receptors (GPCRs) are known as cell-surface receptors mediating approximately two-thirds of human hormones and one-third of Food and Drug Administration (FDA) approved drugs.^1–9^ The GPCRs play a crucial role in the signaling cascade inside a cell making them essential targets in drug design and molecule-based therapeutics,^3,10^ such as in cancer therapy.^9,11^ However, currently, only about 25% of potential druggable GPCRs are being targeted in the drugs due to insufficient experimental and computational knowledge about the mechanism of conformational changes in the receptors.^12^ The GPCRs agonists are remarkably diverse including photons, odorants, viruses, vitamins, peptides, and non-peptidergic hormones.^13–15^ Once a ligand binds to the extracellular part of a receptor, it triggers some conformational changes in the binding site. The sequential conformational changes in amino acids will cause large movements in transmembrane helices.^16–18^ Throughout a series of conformational changes in GPCR, the receptor becomes activated and then messages of the ligand will be relayed to the cell.^19^ The signaling transition is defined as a sequence of conformational changes in the receptor preparing the intracellular region of it to be coupled with the G protein. However, any disruption in the signaling transition in GPCRs may cause some diseases, for example, it can amplify cancer progression.^11,20,21^ Due to the significance of the ligand-mediated signaling pathway, the GPCRs conformational changes have been the subject of remarkable academic efforts to understand diseases associated with the signaling networks. ^12,22,23^ Undesired pathways in the GPCRs may also inhibit the accurate message-passing of the drugs into the cell. That will result in either side effects or low efficacy of the drugs targeting GPCRs. Experimental and theoretical methods (crystallography, NMR, single-molecule force spectroscopy, and spectroscopic methods such as FRET/BRET) are being used to investigate the GPCRs’ signaling mechanism.^24,25^ By investigating the conformational changes in GPCRs within activation states, we will be able to identify the transition pathways connecting the intermediate states with respect to activity levels of the receptor. However, the structural features of intermediate states that coordinate signaling pathways in GPCRs remain poorly understood. It is because of the time resolution required for resolving the dynamics of a protein both experimentally and computationally. Generally, experimental techniques can only provide a few snapshots of GPCR structures (normally active and inactive states) which is insufficient to map the signaling pathways connecting all states of activation. On the other hand, computational methods such as Molecular Dynamics (MD) simulations that can inform about dynamics of the proteins require significant statistical sampling and long trajectories. Obtaining sufficient trajectories for a complete GPCRs activation process via MD simulation takes milliseconds to second,^26–28^ which is computationally expensive. One solution is to run hundreds of short simulations and extract knowledge from trajectory aggregation. Specifically, for GPCRs, multiple experimental crystal structures at different activity levels are available, therefore, different MD simulations could be initiated to collect robust statistics. With tremendous numbers of trajectories, a significant challenge arises from the high-dimensional data and interpretation. To interpret the high-dimensional data, machine learning (ML) models can be used to gain physical insight into the correlation between conformational structures of GPCRs and their activation states.^29–33^ In this study, we develop an ML model based on available experimental information on GPCRs to predict the activity level of a given GPCR structure. Using this model, we evaluate the activity levels of thousands of trajectories of *β*_2_*AR* receptor and predict the transition pathways between states of activation by ordering the activity levels corresponding to protein structural features. This is achieved by combining the reaction coordinates and the estimated activity levels. By taking the advantage of our model, multiple intermediate states that give rise to activity level of the receptor will be identified. In addition, using this method the strength of contribution of each amino acid to activity level can be assessed in order to highlight principal residues driving the transition between activation states in GPCRs. Eventually, for the purpose of enhancing efficacy of drugs, we can design drugs that mainly target the principal amino acids in the receptors.

## Methods

Fig.1 introduces the overall framework of our developed method. The stages of the method are as follows: we first prepared an appropriate experimental dataset of receptors containing protein structure information, state of activation, and activity level corresponding to the conformational structure of the receptors. With this dataset, we defined biophysics-aware features through feature engineering as input to train different shallow ML models.

**Figure 1:**
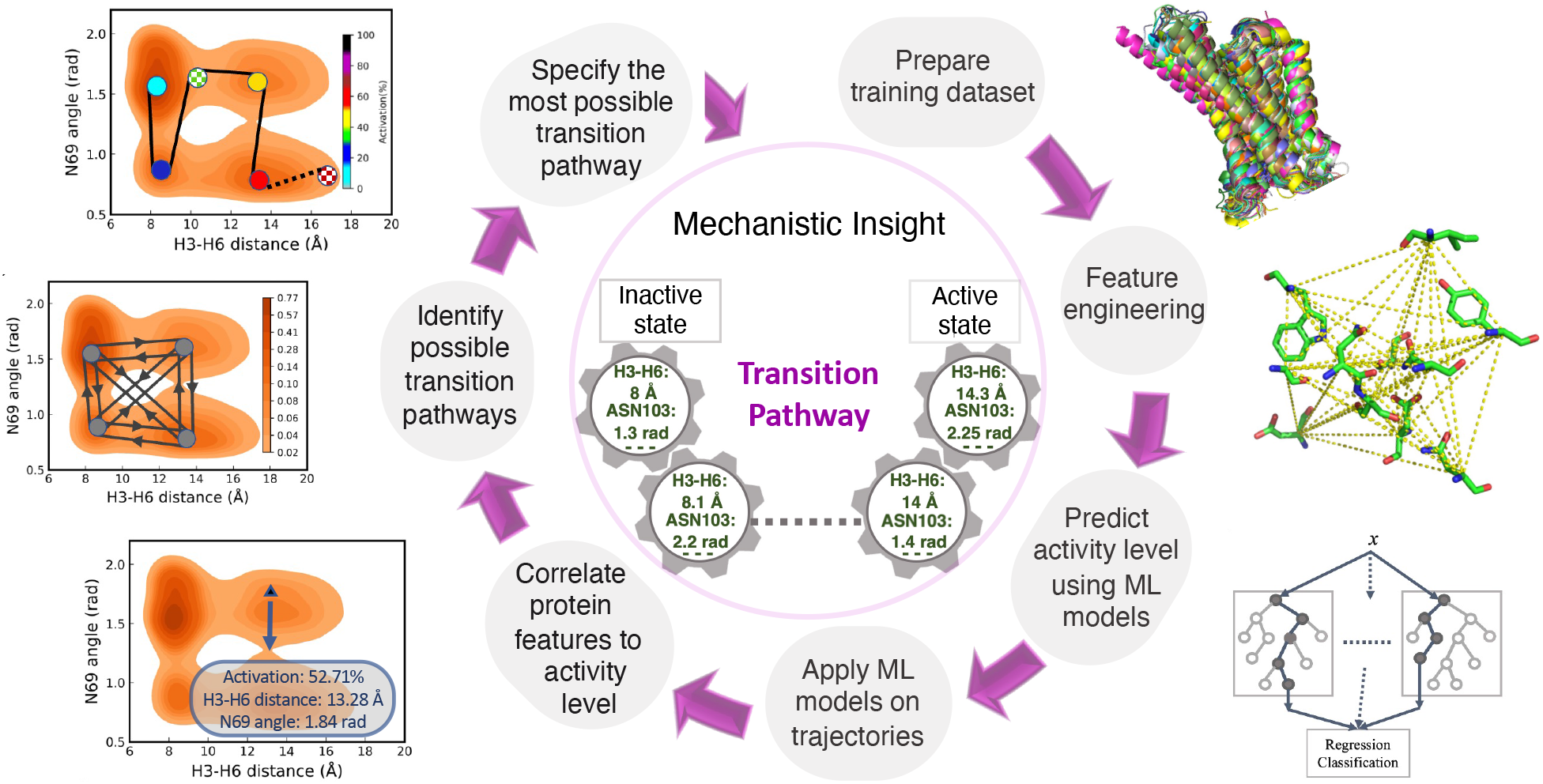
Framework of transition pathway identification between different activation states of a receptor. It consists of training-set preparation, feature engineering, activity level prediction using shallow ML models, applying the most accurate ML model on MD trajectories, correlating protein structural features to the activity level, and finally identifying the most probable transition path between inactive, intermediate, and active states in GPCRs.

### Dataset and preprocessing

The dataset we used for training ML models is taken from GPCRdb database.^34^ It provides experimental information on GPCRs such as ligand-binding constants, phylogenetic diagrams, protein binding, configuration, and signal transduction, as well as different categories of analysis tools and computational data such as homology models and multiple sequence alignments.^34,35^ The training dataset includes 555 protein structures of GPCRs containing 105 unique receptor types^35^(Supporting Information). The PDB database contains threedimensional structural data of proteins and drug molecules.^36^ The state of activation and activity level of the receptors are also obtained from the GPCRdb database.^34^ To prepare the dataset for training ML models, we extract only structures of transmembrane (TM) domains since they play crucial roles in the activation process of GPCRs. The truncated structures of 555 proteins are then spatially aligned to ensure that amino acids located at the same positions are extracted from all the receptors for feature engineering (Supporting Information section 1).

### Feature engineering

With the training dataset, we are able to define a classification task to estimate state of activation and a regression task to predict activity level of a given receptor. In order to develop an ML model, we need to engineer biophysics-aware features (recent knowledge of GPCR structural features to design these features with the goal of achieving higher prediction accuracy) from the dataset to train ML models. Although all GPCRs possess seven-transmembrane structures, only a limited number of amino acids are conserved among them, meaning that similar amino acids are located at the same positions in GPCRs. To train ML models, we included only conserved features suggesting that they may have similar functions in the activation processes.^37^ In this work, we focused on polar network and NPxxY motif contributing to GPCRs signal transduction (Fig.2).^38^ A polar network is a group of amino acids located mainly in the first, second, third, sixth, and seventh transmembrane domains^37^ (Fig.2a,b). The network includes the hydrogen bonds stabilizing the active and inactive states of GPCRs. Comparing active and inactive structures of GPCRs, the polar network has to be rearranged to achieve the active conformation.^37^ In our previous work,^32^ we trained ML models with features comprising of contact distances between two amino acids in random 21-pairs of residues where at least one residue in each pair engaged in the polar network. With that features, ML models could provide activation prediction accuracy of 93.69% for classification task.^32^ However, since the main focus of this study is regression task (i.e activity level prediction), we fine-tuned different combinations of structural features to achieve higher accuracy. We found out that ML models trained with only conserved features can provide higher activation prediction accuracy. Hence, in this study we measured *C_α_* contact distance between each pair of amino acids engaged in the polar network, as shown in Fig.2c (Table S1). Moreover, we computed angular features from the NPxxY motif which is a conserved sequence of amino acids in the seventh transmembrane region^39^ (Fig.2d,e). For this sequence, we defined angle feature between O-C-N atoms in N322^7.49^, P323^7.50^, and Y3 2 6^7.53^ residues involved in NPxxY motif (Fig.2f). In total, we trained ML models with 58 features including 55 contact distances and 3 angle features.

**Figure 2:**
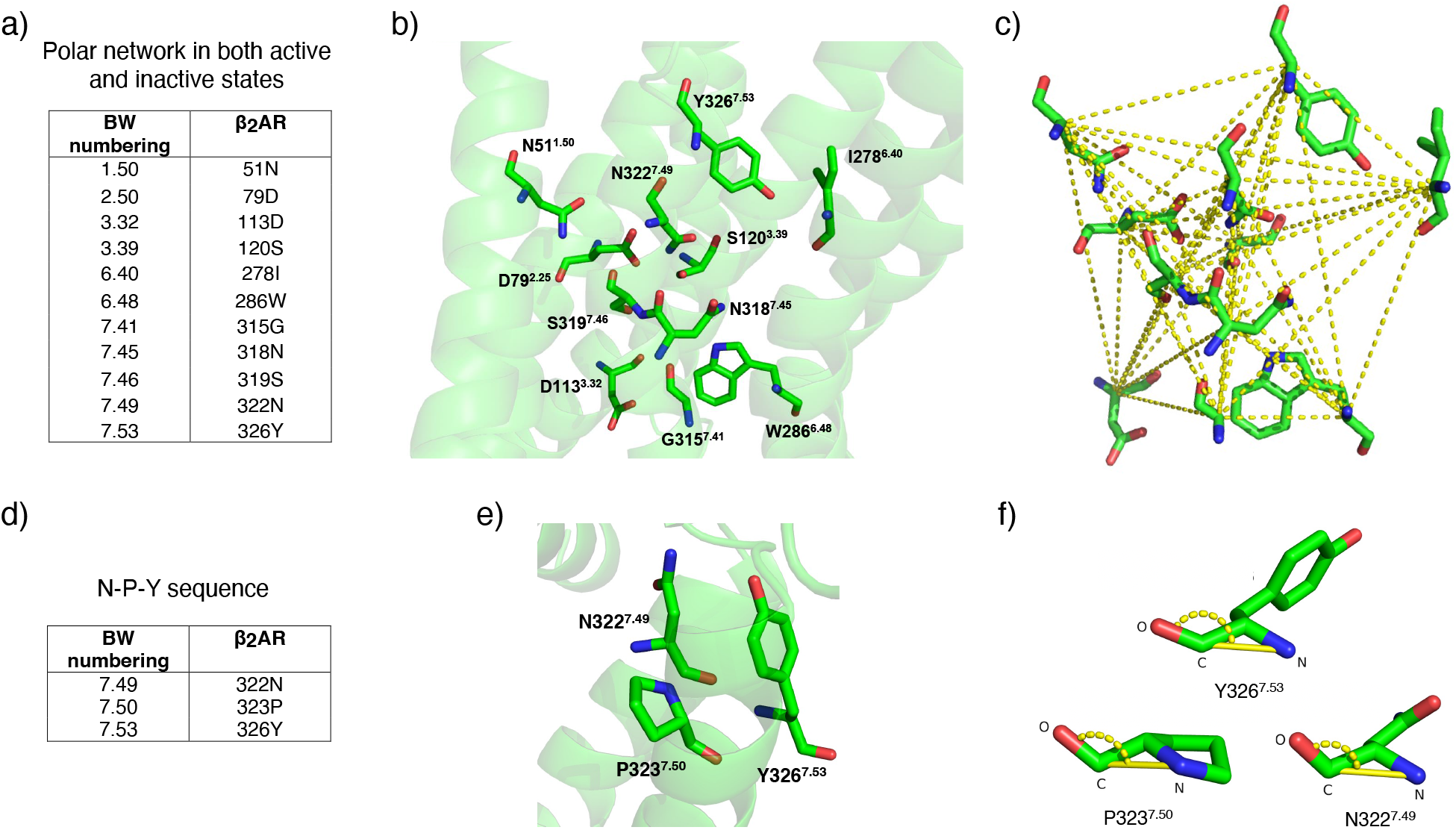
Feature engineering and the type of features selected for activity level prediction using ML. Ballesteros-Weinstein numbers are used to label the amino acids. (a) Amino acids engaged in the polar network in *β*_2_*AR* receptor. (b) Arrangement of polar network residues and their positions in the receptor. (c) Yellow dashed lines represent contact distances between amino acids in the polar network used as input to the ML models. (d) Amino acids involved in the NPxxY motif in the receptor. (e) Position of *N*322^7.49^, *P*323^7.50^, and *Y*326^7.53^ amino acids (as N, P, and Y residues in NPxxY motif) on the receptor. (f) angle features between O-C-N atoms in the N, P, and Y residues used as input to the ML models. Both features are conserved in the active, intermediate, and inactive states of the β_2_AR receptor.

### Structure-activity ML predictor

With respect to the size of the features computed in the previous section and the training dataset, we benchmarked different shallow ML models to predict the activity level of a given receptor. We trained three shallow ML methods including Decision Tree, Random Forest, and XGBoost. Random Forest is an ML method that contains decision trees and output is the mode of trees, while the XGBoost model implements a gradient-boosted trees algorithm to minimize the loss.^40,41^ We used the scikit-learn package to benchmark shallow ML models.^42^ Five-fold cross-validation is performed for these models for both classification and regression tasks. The ML classifiers are then applied to predict the activation state of receptors (in three classes: inactive, intermediate, and active states). The ML regressors are also implemented to predict the activity level of receptors (in the range of 0%-100%).

Table 1 demonstrates performance of the ML models in activation state and activity level prediction. The accuracy and mean absolute error (MAE) for classification and regression tasks are reported, respectively, as two metrics to estimate the performance of the ML models. The results reveal that the XGBoost model predicts both activation state and activity level of a given receptor with higher accuracy compared to the Decision Tree and Random Forest models (Table 1).

**Table 1:** The XGBoost, Random Forest, and Decision Tree model performances in classification and regression tasks. For regression, we reported the mean absolute error (MAE) with standard deviations in parenthesis.

| ML Model | State prediction accuracy (%) | MAE for activation regression (%) |
| --- | --- | --- |
| XGBoost | 97.37 (0.02) | 8.55 (0.67) |
| Random Forest | 94.48 (0.02) | 9.52 (0.81) |
| Decision Tree | 90.45 (0.02) | 18.05 (1.60) |

## Results and discussion

We applied the most accurate ML model (XGBoost) on MD trajectories in order to estimate the correlation between activity level and conformational state (structure) of receptors (for each frame). The main goal, which is to interpret the transition pathway between active, intermediate, and inactive states of GPCRs, is achievable by ordering the activity levels corresponding to the structural properties of the receptors.

### Activity levels of trajectories

MD simulation provides insights into structural ensembles and dynamics of proteins.^43–47^ Using MD trajectories and XGBoost model, we can predict the activity level of each frame of the trajectory and build transition pathways between states of activation by correlating protein structural features to the activity. This can be achieved via statistical sampling and finding stable states. In this study, we used MD simulation trajectories of *β*_2_*AR* receptor.^18^ For the dataset, Google’s Exacycle cloud was used to simulate two milliseconds dynamics of the receptor.^18^ The trajectory dataset is simulated based on three different seeds (initial states) of β2AR receptor including apo; the receptor without any ligand, inverse agonist; with inverse agonist carazolol bound to the receptor, and agonist; with agonist BI-167107 coupled to the receptor.^18^ For each dataset, 5000 MD trajectories (including ~ 150000 frames) of *β*_2_*AR* receptor are randomly chosen (Fig.3). By applying the ML model to the MD trajectories, we are able to interpret how the activity levels of a receptor vary with regard to the dynamics. To apply the XGBoost model on every single frame of the trajectory, we first prepared the input to the ML model (as explained in the feature engineering section) for each frame, and then used the ML model to predict the corresponding activity level. It is necessary to present predicted activity levels of the trajectories in terms of a known structural feature of GPCRs. One of the experimentally best-known features of GPCRs related to activation states is Helix3-Helix6 (H3-H6) distance. It is notable that the H3-H6 distance in the inactive and active structures of *β*_2_*AR* receptor is ~ 8.4Å and 14. 1Å, respectively. To investigate how the activity levels change in terms of a protein structural feature, we plotted related density histogram in different conformational states of the receptor mapped on corresponding predicted activity levels (Fig.3). For the density histograms in Fig.3 we measured the H3-H6 distance as *C_α_* contact distance between *R*131^3.50^-L272^6.34^ amino acids at each frame of the simulations. On the histograms, each bar is colored based on the average of predicted activities of data points associated with it. All figures showing the density histogram generated by H3-H6 distance contain two points (shown by triangle features) introducing the experimental H3-H6 distance in the active and inactive crystal structures of *β*_2_*AR* receptor. These two points are used to validate the performance of our model. With respect to these two points, we observe that the ML model predicted inactive, active, and intermediate states mostly around ~ 8.4Å, ~ 14.1Å, and between ~ 8. 4A-14.1Å, respectively. This result reveals that the ML predictor is accurate enough for both the classification and regression tasks. In addition, using Fig.3b containing the inverse agonist dataset of *β*_2_*AR* receptor, we investigated the relationship between H3-H6 distance and the activity level of the receptor. Fig. S2 shows that in the inactive state, the activity level is correlated to the H3-H6 distance much stronger than in the active or intermediate states, meaning that when the receptor is in the inactive state a very small change in H3-H6 distance makes a large change in its activity level.

**Figure 3:**
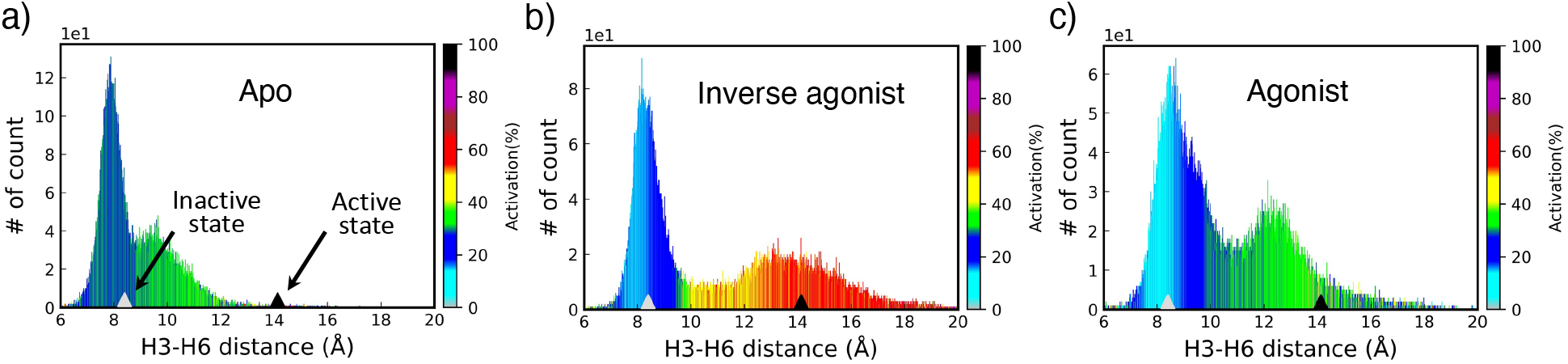
Histograms of H3-H6 distance for different conformational states of *β*_2_*AR* receptor. The colorbars show the activity levels (%) predicted by the XGBoost model. The parameter used for generating the histograms is *C_α_* contact distance between *R*131^3.50^-L272^6.34^ amino acids. (a) Apo; the receptor without any ligand. (b) Inverse agonist; with inverse agonist carazolol bound to the receptor. (c) Agonist; with agonist BI-167107 coupled to the receptor.

### Identifying the most probable transition pathway

In order to identify the complete transition pathway between states of activation, the correlations between the activity level and all structural features of a receptor (for example, angular changes in each amino acid or residue pair-wise distances) are essentially required. This will provide an n-dimensional density landscape mapped on predicted activity levels. To be able to visualize such a correlation, we projected 2D density maps using two structural features of *β*_2_*AR* receptor in the inverse agonist dataset. For all the 2D plots, one dimension (on the horizontal axis) is H3-H6 distance. Fig.4a represents the histogram of *S*329^8.47^ angle feature and its combination with the histogram of H3-H6 distance provides a density landscape as shown in Fig.4b. Fig.4c,d illustrate the density projection of *N*69^2.40^ angle feature and density landscape corresponding to the feature and H3-H6 distance. Fig.4b,d also introduce the average activity levels in each distinct region in the landscapes separated by dashed lines. The density landscapes verify that the distribution of densities varies with respect to the protein features and activity levels, meaning that each region of density has its own majority of activation states. For example, Fig.4b shows that going from one peak of density at H3-H6 distance ~ 8Å to another peak at ~ 14Å both of the average of activity level and *S*329^8.47^ angle feature enhance (average of activity level increases from 19.29% to 54.81% and *S*329^8.47^ angle feature from 1.8 rad to 2.2 rad). However, such changes are more complicated in the density landscapes including *N*69^2.40^ angle feature and H3-H6 distance, as shown in Fig.4d. It can be shown in Fig.5a,d that without using ML models, there are multiple potential pathways connecting the peaks of densities (shown with grey circles) in different directions (shown with arrows). We used three steps to find the most probable pathway: 1. find all the high-density peaks over the landscape and connect them (two by two) through the minimum-density paths (Fig.5a,d), 2. predict the average activity levels all over the landscape. Here we measured the average activity levels in small areas 3 × 10^-3^ rad.A (Fig.5b,e), and 3. connect the states in ascending order of activation in order to build the transition pathway from inactive to active state. (Fig.5c,f). Therefore, the combination of density of states and the activity levels can yield the most probable pathways between activation states of a receptor. Note that for visualization purposes we performed transition pathway identification only for two features. However, it should be done in high dimensional space (n-dimension corresponding to n-structural features of the receptor). Then, we can identify the global most possible transition pathway between activation states of the receptor by ranking all states in terms of activity levels. Fig.5g introduces a few pieces of a possible transition pathway in terms of H3-H6 distance, *S*329^8.47^ angle feature, and *N*69^2.40^ angle feature. It can be observed in Fig.5 that H3-H6 distance and *S*329^8.47^ angle feature are increasing as the activation increases, while the *N*69^2.40^ angle feature is oscillating for similar changes in activation. Therefore, our framework not only is capable of identifying the path of transition between activation states but also can evaluate the strength of correlation between each protein structural feature to activity level of the receptor. This provides significant mechanistic insight into GPCRs activation and drug discovery. By estimating the contribution of each amino acid in the activation process, we can design drugs that mainly target the principal amino acids driving the transition pathway between activation states, in order to minimize side effects or maximize efficacy of the drugs targeting GPCRs.

**Figure 4:**
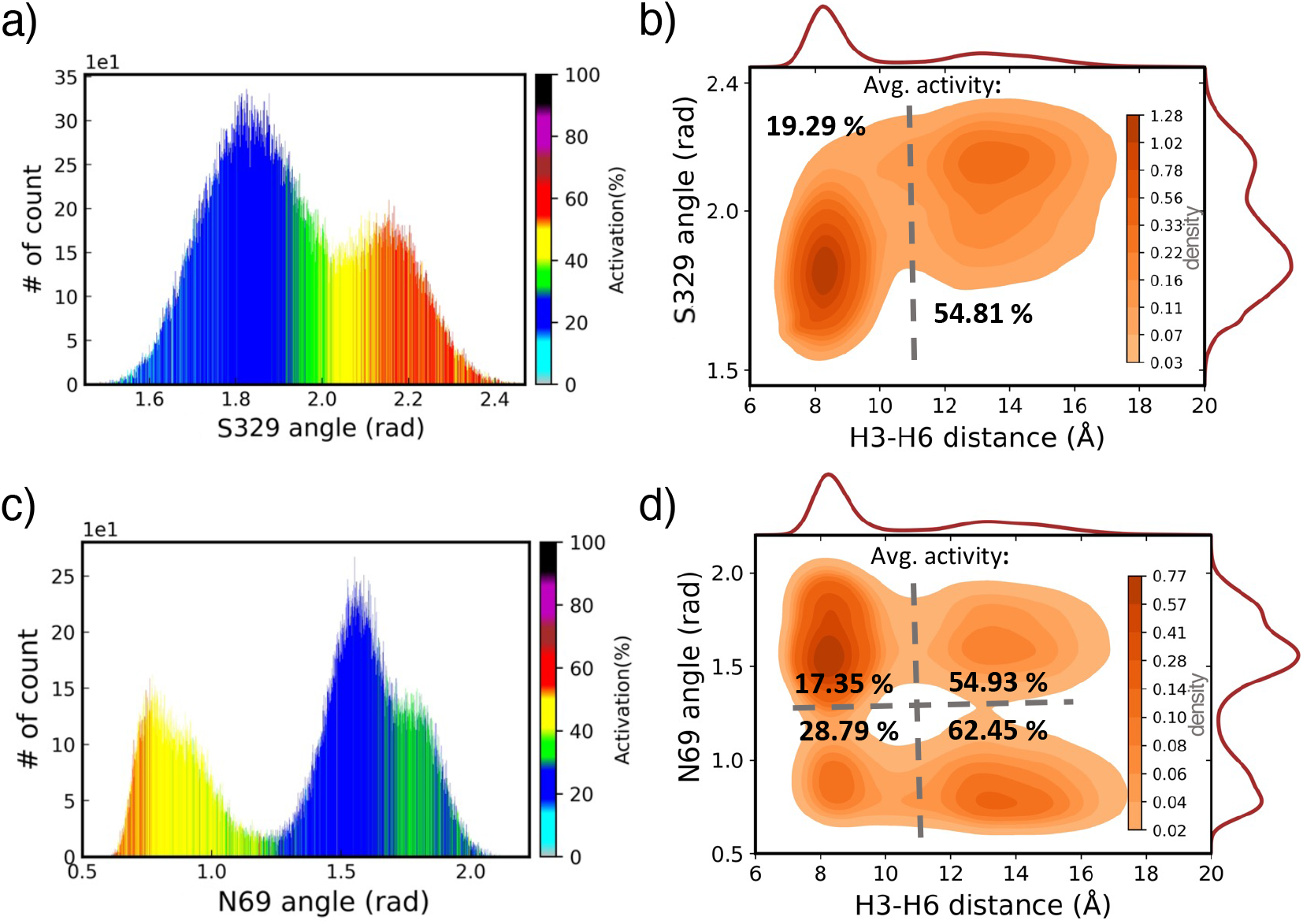
1D and 2D density histograms corresponding to activity level predictions in the inverse agonist dataset of *β*_2_*AR* receptor projected on two reaction coordinate features. (a) 1D density histogram projected on *S*329^8.47^ residue angle feature. (b) Density landscape of *S*329^8.47^ residue angle feature vs H3-H6 distance with average of activation denoted for two regions separated by the dashed line. (c) 1D density histogram with predicted activity levels generated by angle feature of N69^2.40^ residue. (d) Density landscape of N69^2.40^ residue angle feature vs H3-H6 distance with average of activation computed for four regions separated by the dashed line. The colorbars in (a) and (c) represent the activity level predicted by the XGBoost model and the colorbars in (b) and (d) show the density of states.

**Figure 5:**
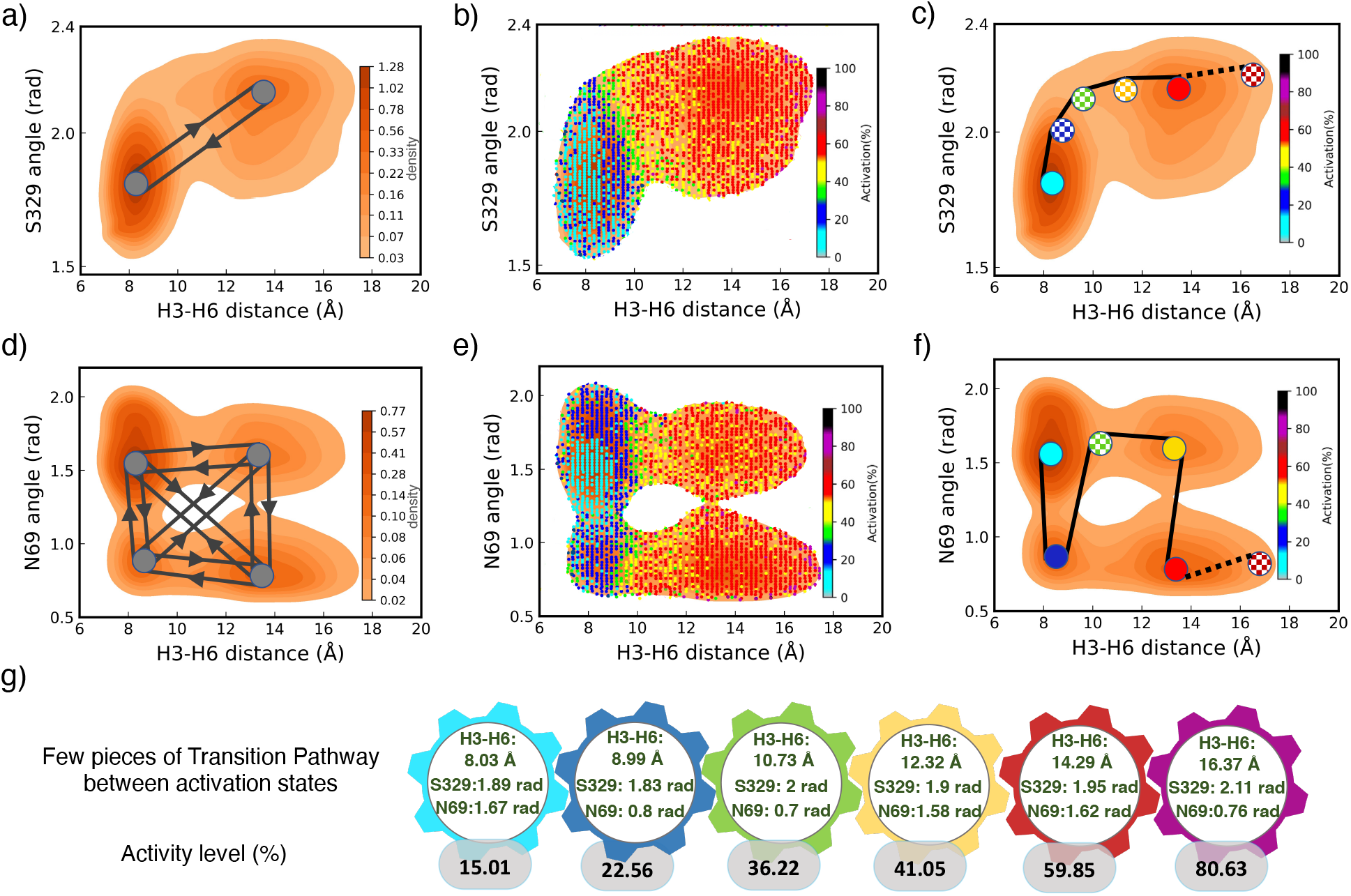
Density landscape of *β*_2_*AR* receptor projected on H3-H6 distance and angle features with activity levels. (a) Density landscape of *S*329^8.47^ angle feature vs H3-H6 distance with two possible pathways based on the density peaks. (b) Predicted activity levels on density landscape of *S*329^8.47^ angle feature and H3-H6 distance at small areas (3 × 10^-3^ rad.Å). (c) The proposed activation pathway for the two features (*S*329^8.47^ angle feature and H3-H6 distance) with a gradual change in the activity levels shown by filled and patterned circles. (d) Few potential transition pathways between four states populated with density peaks of *N*69^2.40^ angle feature and H3-H6 distance. (e) Predicted activity levels for density landscape of *N*69^2.40^ angle feature and H3-H6 distance at the small areas (3 × 10^-3^ rad.A). (f) The proposed activation pathway for the two features (*N*69^2.40^ angle feature and H3-H6 distance) with a gradual change in the activity levels shown by the filled and patterned circles. (g) A few of three feature statuses (H3-H6 distance, *S*329^8.47^ angle feature, and *N*69^2.40^ angle feature) ranked in ascending order of activity level. The filled circles (in (c) and (f)) present the states recognized by the high-density peaks while the patterned filled circles are samples showing the activity levels on the ways between the density peaks.

## Conclusion

In this work, we have developed an ML-based framework to predict the most probable transition pathway between activation states of GPCRs based on structural features of the receptors. To achieve this, we devised an ML model to predict the activity level of GPCRs. We considered two conserved biophysics-aware feature types containing the polar network and NPxxY motif to train shallow ML models. The XGBoost model can be used to predict activation state and activity level of GPCRs with 97.37% accuracy and 8.55% MAE, respectively. We then applied the ML model to MD trajectories of *β*_2_*AR* receptor to correlate protein structural dynamics to the activity levels with the goal of finding the most probable transition pathway between states of activation. By combining density of states and the activity levels, we were able to find the most probable pathway. In this study, we featurized two residues (for easy visualization) in all the trajectories and then projected their density landscapes providing many possible transition pathways between density peaks. To identify the most probable pathway, we evaluated the activity levels in all the states over the landscape, and then ranked the activity levels correlated to the defined structural features in ascending order to identify the path of transition from inactive to active states. The complete transition pathway can be predicted by considering all n-structural features of the receptor. Overall, this framework provides valuable knowledge on the activation mechanism of GPCRs and can significantly accelerate the discovery of drugs targeting GPCRs.

## Acknowledgement

The authors gratefully acknowledge the use of MD simulation trajectories of *β*2*AR* receptor dataset provided by Stanford University.^18^ We thank Parakarsh Yadav for his help with data preprocessing. This work is supported by the Center for Machine Learning in Health (CMLH) at Carnegie Mellon University and a start-up fund from Mechanical Engineering Department at CMU.

