## Supporting Information for "Activity Map and Transition Pathways of G Protein Coupled Receptor Revealed by Machine Learning"

#### Contents

**1 GPCRs structure data preprocessing** **2**

**2 Residue-pairs in the polar network** **2**

**3 Activation rates in transition between states of $\beta_2 AR$ receptor** **3**

**4 Dataset organization** **4**

**References** **4**

### 1 GPCRs structure data preprocessing

The training dataset contains 555 proteins from the RCSB server. First, we aligned all these proteins since the features we defined for training ML models are position-dependent and include contact distances of residues engaged in the polar network and angle features of residues involved in the NPxxY motif. To overcome the challenge of feature extraction, we spatially aligned all the GPCRs to a reference receptor (Fig.S1c). The reference receptor is the Neurotensin Receptor1 (NTSR1, PDB:6UP7) which belongs to Class-A (Rhodopsin) GPCR. All the proteins are aligned by using the align tool of PyMOL software.[1,2] For this study, we only selected amino acids that are on the helical structures and removed all the intracellular loops. This ensures that only the transmembrane domains having significant roles in the activation process are used for extracting features to train ML models.

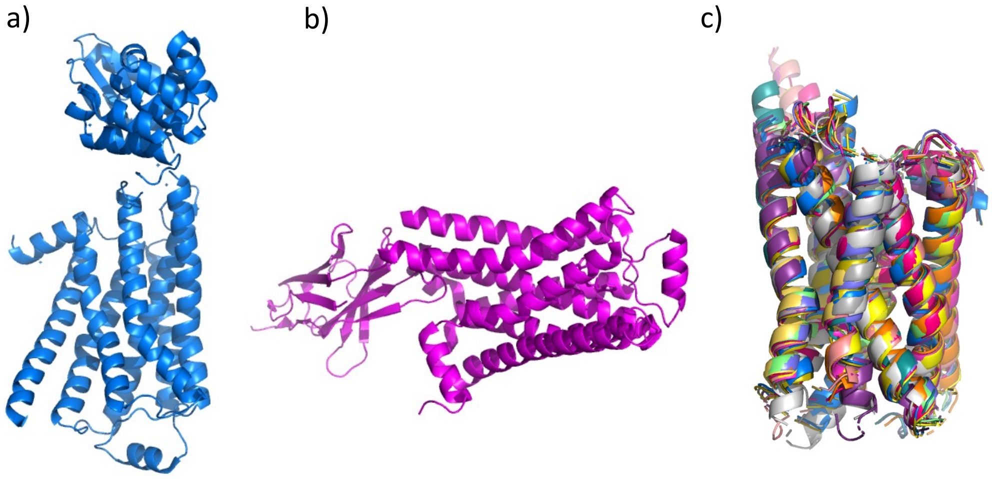

Figure S1: The visualization of the original (a) inactive and (b) active structures of $\beta_2 AR$ receptor along with TM region and other sub-units. (c) Some transmembrane domains of GPCRs structures in the training dataset aligned with the NTSR1 receptor (PDB: 6UP7).

### 2 Residue-pairs in the polar network

Table 1 represents all 55 pairs of residues in the polar network of $\beta_2 AR$ receptor. The $C_\alpha$ contact distance between each of the residue-pairs (shown in Table 1) is measured as input to the ML models.[3] This list is conserved in the active, intermediate, and inactive conformations of $\beta_2 AR$ receptor.

Table 1: List of 55 residue-pairs in polar network of $\beta_2 AR$ receptor. The $C_\alpha$ contact distance between each residue pair in this list is measured to train ML models.

| | | | |
|---|---|---|---|
| $N51^{1.50}$-$D79^{2.50}$ | $N51^{1.50}$-$D113^{3.32}$ | $N51^{1.50}$-$S120^{3.39}$ | $N51^{1.50}$-$I278^{6.40}$ |
| $N51^{1.50}$-$G315^{7.41}$ | $N51^{1.50}$-$N318^{7.45}$ | $N51^{1.50}$-$S319^{7.46}$ | $N51^{1.50}$-$N322^{7.49}$ |
| $D79^{2.50}$-$D113^{3.32}$ | $D79^{2.50}$-$S120^{3.39}$ | $D79^{2.50}$-$I278^{6.40}$ | $D79^{2.50}$-$W286^{6.48}$ |
| $D79^{2.50}$-$N318^{7.45}$ | $D79^{2.50}$-$S319^{7.46}$ | $D79^{2.50}$-$N322^{7.49}$ | $D79^{2.50}$-$Y326^{7.53}$ |
| $D113^{3.32}$-$I278^{6.40}$ | $D113^{3.32}$-$W286^{6.48}$ | $D113^{3.32}$-$G315^{7.41}$ | $D113^{3.32}$-$N318^{7.45}$ |
| $D113^{3.32}$-$N322^{7.49}$ | $D113^{3.32}$-$Y326^{7.53}$ | $S120^{3.39}$-$I278^{6.40}$ | $S120^{3.39}$-$W286^{6.48}$ |
| $S120^{3.39}$-$N318^{7.45}$ | $S120^{3.39}$-$S319^{7.46}$ | $S120^{3.39}$-$N322^{7.49}$ | $S120^{3.39}$-$Y326^{7.53}$ |
| $I278^{6.40}$-$G315^{7.41}$ | $I278^{6.40}$-$N318^{7.45}$ | $I278^{6.40}$-$S319^{7.46}$ | $I278^{6.40}$-$N322^{7.49}$ |
| $W286^{6.48}$-$G315^{7.41}$ | $W286^{6.48}$-$N318^{7.45}$ | $W286^{6.48}$-$S319^{7.46}$ | $W286^{6.48}$-$N322^{7.49}$ |
| $G315^{7.41}$-$N318^{7.45}$ | $G315^{7.41}$-$S319^{7.46}$ | $G315^{7.41}$-$N322^{7.49}$ | $G315^{7.41}$-$Y326^{7.53}$ |
| $N318^{7.45}$-$N322^{7.49}$ | $N318^{7.45}$-$Y326^{7.53}$ | $S319^{7.46}$-$N322^{7.49}$ | $S319^{7.46}$-$Y326^{7.53}$ |
| $N51^{1.50}$-$W286^{6.48}$ | $N51^{1.50}$-$Y326^{7.53}$ | $D79^{2.50}$-$G315^{7.41}$ | $D113^{3.32}$-$S120^{3.39}$ |
| $D113^{3.32}$-$S319^{7.46}$ | $S120^{3.39}$-$G315^{7.41}$ | $I278^{6.40}$-$W286^{6.48}$ | $I278^{6.40}$-$Y326^{7.53}$ |
| $W286^{6.48}$-$Y326^{7.53}$ | $N318^{7.45}$-$S319^{7.46}$ | $N322^{7.49}$-$Y326^{7.53}$ | |

### 3 Activation rates in transition between states of $\beta_2 AR$ receptor

Fig.S2 shows the histogram of $\beta_2 AR$ receptor generated by H3-H6 distance for the inverse agonist dataset mapped on activity levels. The piece-wise linear slopes illustrate how the activity levels change with respect to the H3-H6 distance. As shown in the plot, the slope of activity level in terms of H3-H6 distance in the inactive states is much higher compared to the active or intermediate states.

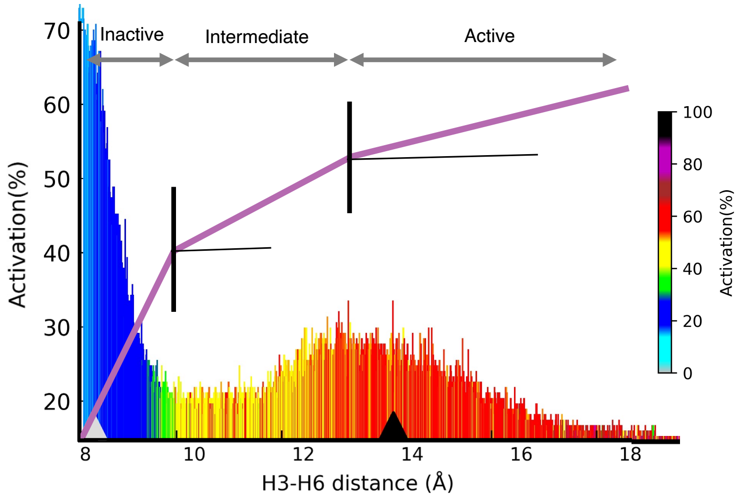


Figure S2: Different rates of activity with respect to H3-H6 distance in the inverse agonist dataset of $\beta_2AR$ receptor. The black triangle on the x-axis represents the H3-H6 distance in the crystal active structure of $\beta_2AR$ receptor and the grey one shows the H3-H6 distance in the crystal inactive structure of the receptor. The colorbar shows the predicted activity levels by XGBoost model (%0-%100)

### 4 Dataset organization

A zip file containing the processed PDB structures of the GPCRs, information on the classification and regression labels of the receptors, and all the scripts for ML models used in this study are available here: https://github.com/BaratiLab/GPCRpath
